## Supplemental Figures S1-S3 and Tables 1-4 for "Peripheral opioid receptor antagonism alleviates fentanyl-induced cardiorespiratory depression and is devoid of aversive behavior"

Affiliations:

**The file includes:**

Figures S1 to S3, Tables 1-4

1071

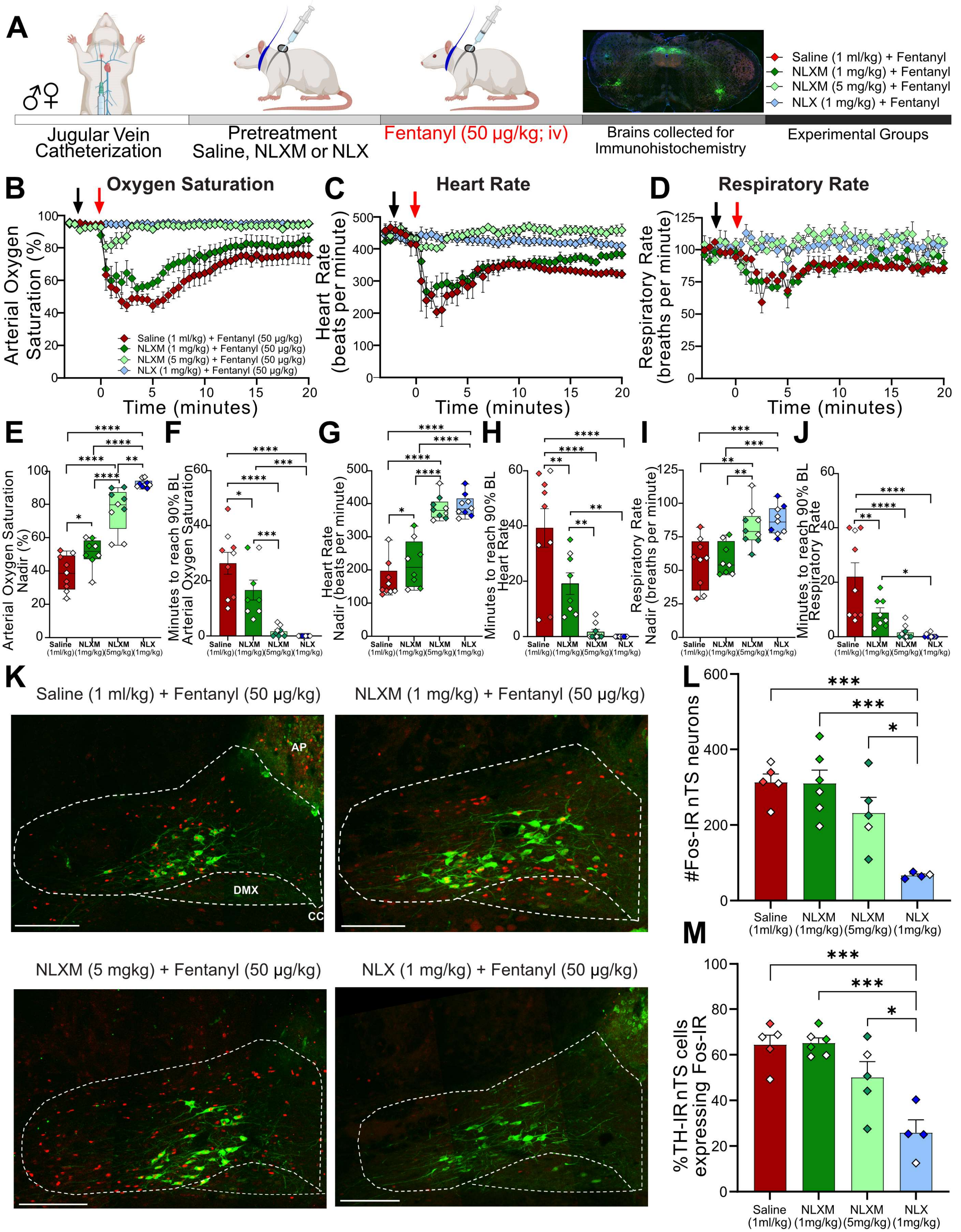
A
Jugular Vein Catheterization
Pretreatment Saline, NLXM or NLX
Fentanyl (50 μg/kg; iv)
Brains collected for Immunohistochemistry
Experimental Groups
Saline (1 ml/kg) + Fentanyl
NLXM (1 mg/kg) + Fentanyl
NLXM (5 mg/kg) + Fentanyl
NLX (1 mg/kg) + Fentanyl
B Oxygen Saturation
Arterial Oxygen Saturation (%)
Time (minutes)
Saline (1 ml/kg) + Fentanyl (50 μg/kg)
NLXM (1 mg/kg) + Fentanyl (50 μg/kg)
NLXM (5 mg/kg) + Fentanyl (50 μg/kg)
NLX (1 mg/kg) + Fentanyl (50 μg/kg)
C Heart Rate
Heart Rate (beats per minute)
Time (minutes)
D Respiratory Rate
Respiratory Rate (breaths per minute)
Time (minutes)
E
Arterial Oxygen Saturation Nadir (%)
F
Minutes to reach 90% BL Arterial Oxygen Saturation
G
Heart Rate Nadir (beats per minute)
H
Minutes to reach 90% BL Heart Rate
I
Respiratory Rate Nadir (breaths per minute)
J
Minutes to reach 90% BL Respiratory Rate
Saline (1ml/kg) NLXM (1mg/kg) NLXM (5mg/kg) NLX (1mg/kg)
K
Saline (1 ml/kg) + Fentanyl (50 μg/kg)
NLXM (1 mg/kg) + Fentanyl (50 μg/kg)
AP
DMX
CC
NLXM (5 mgkg) + Fentanyl (50 μg/kg)
NLX (1 mg/kg) + Fentanyl (50 μg/kg)
L
#Fos-IR nTS neurons
M
%TH-IR nTS cells expressing Fos-IR
Saline (1ml/kg) NLXM (1mg/kg) NLXM (5mg/kg) NLX (1mg/kg)

**Supplemental Figure 1: NLXM attenuates cardiorespiratory depression induced by a higher dose of fentanyl. A**. Schematic showing the timeline for evaluation of reversal of OIRD. Male and Female SD rats underwent jugular catheterization surgery. On the day of the experiment, rats were administered saline (1 ml/kg), NLX (1 mg/kg) or NLXM (1 or 5 mg/kg) intravenously. Two minutes later, rats received intravenous fentanyl (50 µg/kg). Cardiorespiratory parameters were measured for up to 60 minutes after fentanyl administration. The first 20 minutes of time course data showing oxygen saturation (**B**), heart rate (**C**) and respiratory rate (**D**) measurements is shown for all groups. **E-J** shows the nadir of each cardiorespiratory parameter (**E,G,I**) and time to return to 90% of baseline values (pre-saline/fentanyl; **F,H,J**). In all graphs, males are represented by filled symbols, and females by open symbols. For graphs displaying mean nadir values: Oxygen Saturation: One-way ANOVA, $F_{(3,31)}$ = 58.04, p <0.0001;Tukey's post-hoc test, **** p<0.0001; ** p<0.01; *p<0.05; Heart Rate: One-way ANOVA, $F_{(3,31)}$ = 46.34, p <0.0001; Tukey's post-hoc test, **** p<0.0001;  *p<0.05; Respiratory Rate: One-way ANOVA, $F_{(3,31)}$ = 11.12, p <0.001;Tukey's post-hoc test, ***p<0.001, **p<0.01). For graphs displaying time to reach 90% baseline: Oxygen Saturation: (One-way ANOVA, $F_{(3,31)}$ = 23.00, p <0.0001; Tukey's post-hoc test ****p<0.0001; ***p<0.001; *p<0.05; Heart rate: (One-way ANOVA, $F_{(3,31)}$ = 21.13, p <0.0001; Tukey's post-hoc test ****p<0.0001; **p<0.01); Respiratory rate: (One-way ANOVA, $F_{(3,31)}$ = 12.79, p <0.0001; Tukey's post-hoc test, ****p<0.0001; **p<0.01; *p<0.05). **K**. Merged photomicrographs of a coronal brainstem section displaying Fos- and TH- immunoreactivity in the nTS from all groups. Scale bar = 200 µm. **L**. The number of Fos-IR cells was similar between saline and both NLXM treated groups. There were significantly fewer Fos cells in 1 mg/kg NLX-pretreated animals. One-way ANOVA, $F_{(3,16)}$ = 11.40, p <0.001; Tukey's post-hoc ***p<0.001; *p<0.05. **M** shows the percentage of TH-IR nTS neurons expressing Fos-IR. There was no significant difference between saline- and NLXM-pretreated rats. NLX-pretreated rats displayed significantly lower percentage of TH-IR nTS neurons expressing Fos-IR. One-way ANOVA, $F_{(3,16)}$ = 13.06, p=0.0001; Tukey's post-hoc *** p<0.001; * p<0.05).

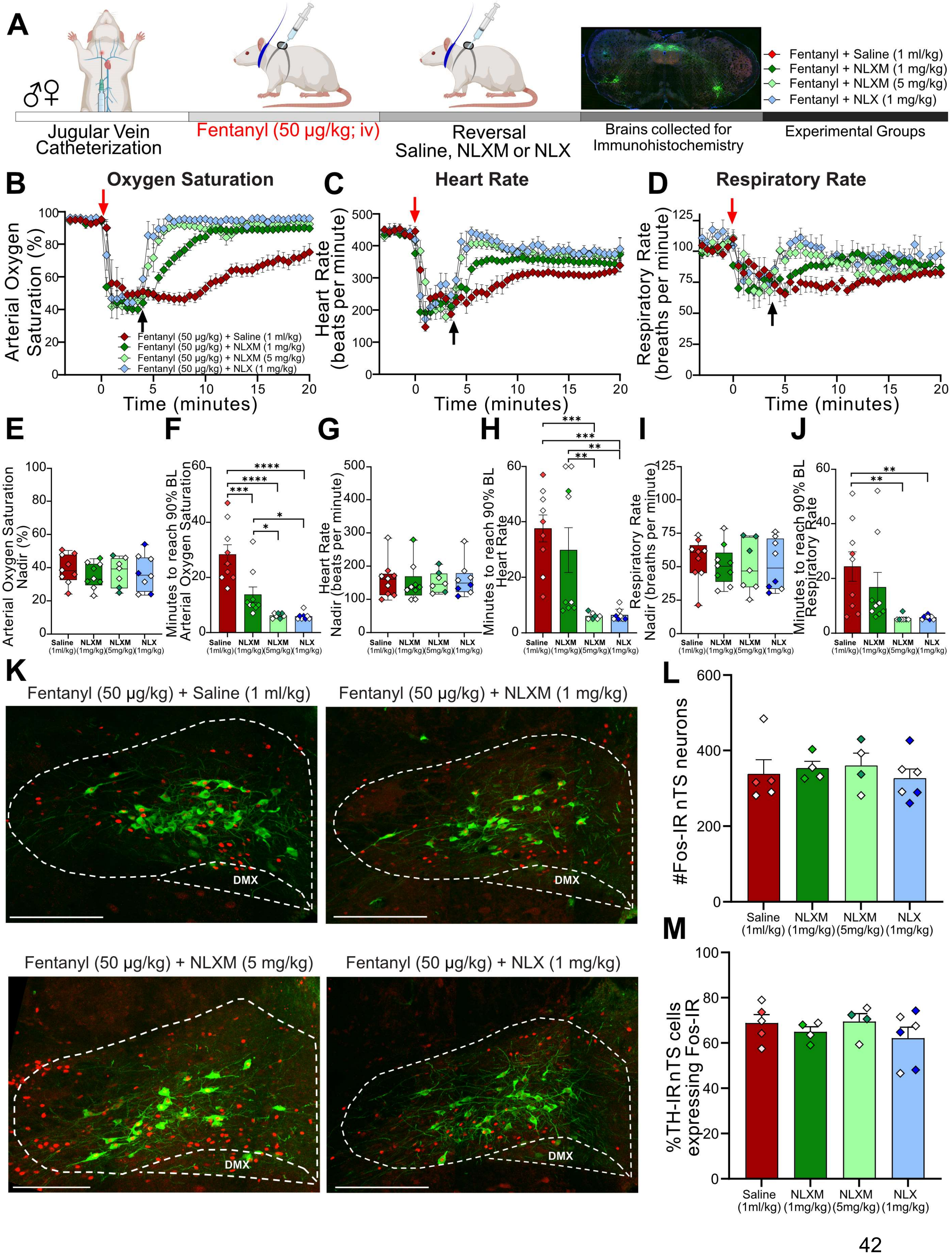

A
Jugular Vein Catheterization
Fentanyl (50 µg/kg; iv)
Reversal Saline, NLXM or NLX
Brains collected for Immunohistochemistry
Experimental Groups
Fentanyl + Saline (1 ml/kg)
Fentanyl + NLXM (1 mg/kg)
Fentanyl + NLXM (5 mg/kg)
Fentanyl + NLX (1 mg/kg)
B
Oxygen Saturation
Arterial Oxygen Saturation (%)
Time (minutes)
Fentanyl (50 µg/kg) + Saline (1 ml/kg)
Fentanyl (50 µg/kg) + NLXM (1 mg/kg)
Fentanyl (50 µg/kg) + NLXM (5 mg/kg)
Fentanyl (50 µg/kg) + NLX (1 mg/kg)
C
Heart Rate
Heart Rate (beats per minute)
Time (minutes)
D
Respiratory Rate
Respiratory Rate (breaths per minute)
Time (minutes)
E
Arterial Oxygen Saturation Nadir (%)
F
Minutes to reach 90% BL Arterial Oxygen Saturation
G
Heart Rate Nadir (beats per minute)
H
Minutes to reach 90% BL Heart Rate
I
Respiratory Rate Nadir (breaths per minute)
J
Minutes to reach 90% BL Respiratory Rate
Saline (1ml/kg) NLXM (1mg/kg) NLXM (5mg/kg) NLX (1mg/kg)
K
Fentanyl (50 µg/kg) + Saline (1 ml/kg)
Fentanyl (50 µg/kg) + NLXM (1 mg/kg)
Fentanyl (50 µg/kg) + NLXM (5 mg/kg)
Fentanyl (50 µg/kg) + NLX (1 mg/kg)
DMX
L
#Fos-IR nTS neurons
M
%TH-IR nTS cells expressing Fos-IR
Saline (1ml/kg) NLXM (1mg/kg) NLXM (5mg/kg) NLX (1mg/kg)

**Supplemental Figure 2: NLXM reverses cardiorespiratory depression induced by a higher dose of fentanyl. A**. Schematic showing the timeline for evaluation of reversal of OIRD. Male and Female SD rats underwent jugular catheterization surgery. On the day of the experiment, rats received intravenous fentanyl (50 µg/kg). Four minutes later, rats received saline (1 ml/kg), NLX (1 mg/kg) or NLXM (1 or 5 mg/kg). Cardiorespiratory parameters were collected for up to 60 minutes after saline or antagonist administration. The first 20 minutes of time course data showing oxygen saturation (**B**), heart rate (**C**) and respiratory rate (**D**) measurements is shown for all groups. **E-J** shows each cardiorespiratory parameter at their respective nadir (**E,G,I**) and time to recover to 90% of baseline (pre-fentanyl; **F,H,J**) administration. In all graphs, males are represented by filled symbols, and females by open symbols. Mean nadir values were assessed in all groups. Oxygen Saturation: One-way ANOVA, $F_{(3,29)} = 0.2892$, $p = 0.8325$; Heart Rate: One-way ANOVA, $F_{(3,29)} = 0.09684$, $p=0.9612$; Respiratory Rate: One-way ANOVA, $F_{(3,29)} = 0.1443$, $p=0.9325$. Mean time to reach 90% baseline was assessed for all parameters. Oxygen Saturation: (One-way ANOVA, $F_{(3,29)} = 18.25$, $p <0.0001$; Tukey's post-hoc test ****p<0.0001; ***p<0.001; *p<0.05; Heart rate: (One-way ANOVA, $F_{(3,29)} = 9.705$, $p=0.001$; Tukey's post-hoc test ***p<0.001; **p<0.01; Respiratory rate: (One-way ANOVA, $F_{(3,29)} = 4.799$, $p <0.01$; Tukey's post-hoc test, **p<0.01). **K**. Merged photomicrographs of a coronal brainstem section displaying Fos- and TH- immunoreactivity in the nTS from all groups. Scale bar = 200 µm. **L**. Neuronal activation was assessed in saline-, NLX and NLXM-treated rats. Number of Fos-IR cells: One-way ANOVA, $F_{(3,15)} = 0.2689$, $p=0.8468$; **M**. Percentage of TH-IR nTS cells expressing Fos-IR: (One-way ANOVA, $F_{(3,15)} = 0.7616$, $p=0.5330$.

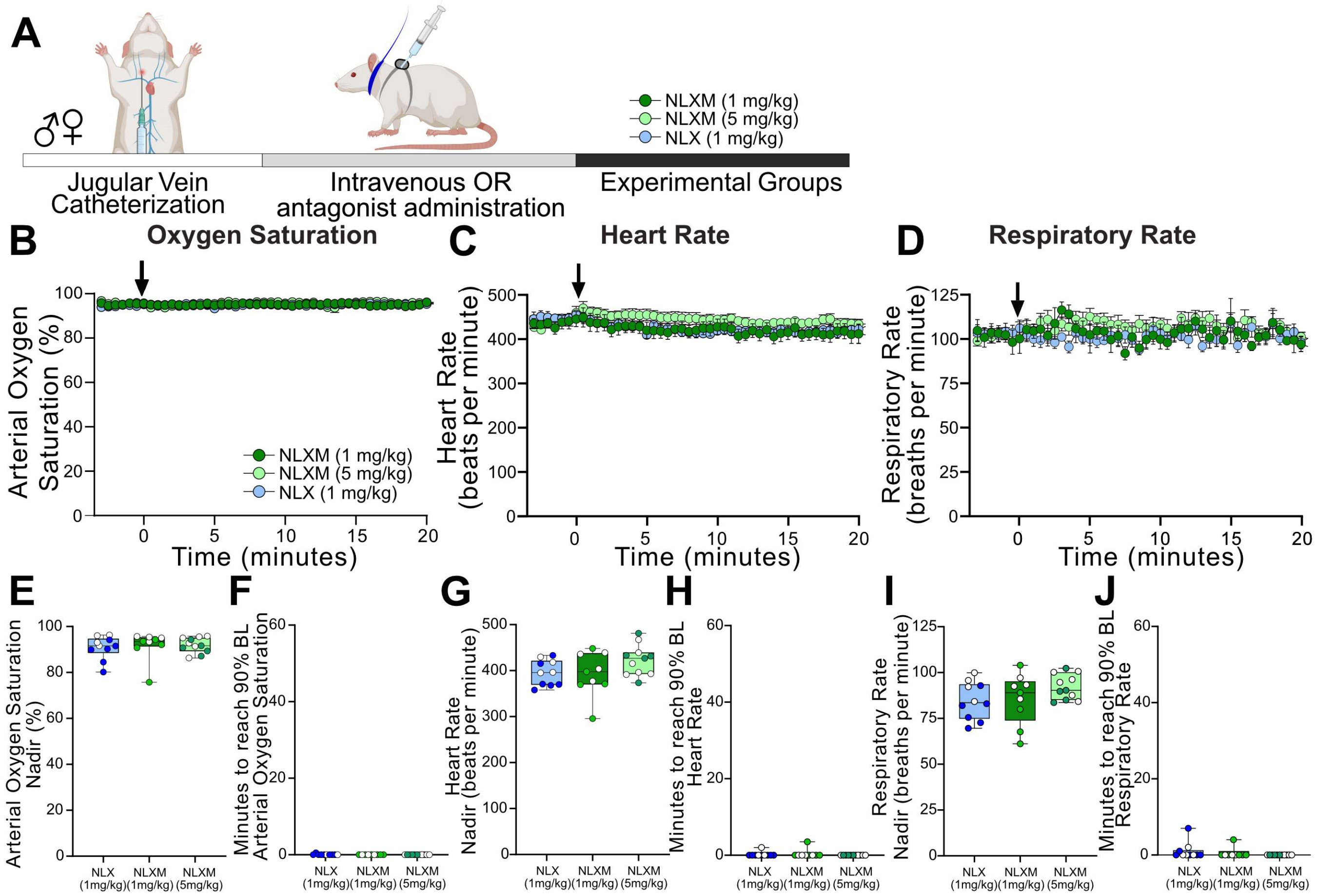
A
Jugular Vein Catheterization
Intravenous OR antagonist administration
Experimental Groups
NLXM (1 mg/kg)
NLXM (5 mg/kg)
NLX (1 mg/kg)
B
Oxygen Saturation
Arterial Oxygen Saturation (%)
Time (minutes)
C
Heart Rate
Heart Rate (beats per minute)
Time (minutes)
D
Respiratory Rate
Respiratory Rate (breaths per minute)
Time (minutes)
E
Arterial Oxygen Saturation Nadir (%)
F
Minutes to reach 90% BL Arterial Oxygen Saturation
G
Heart Rate Nadir (beats per minute)
H
Minutes to reach 90% BL Heart Rate
I
Respiratory Rate Nadir (breaths per minute)
J
Minutes to reach 90% BL Respiratory Rate
NLX (1mg/kg)
NLXM (1mg/kg)
NLXM (5mg/kg)

**Supplemental Figure 3: Opioid Receptor antagonism has no effect on baseline cardiorespiratory parameters in drug-naïve rats. A**. Schematic showing the timeline for evaluation of cardiorespiratory parameters following administration of opioid receptor antagonists in catheterized male and female SD rats. **B-D.** Rats received either intravenous (black arrow) 1 mg/kg NLX (blue symbols), 1 mg/kg NLXM (light green symbols) or 5 mg/kg NLXM (dark green symbols). Cardiorespiratory parameters were measured up to 20 minutes. The nadir and recovery of oxygen saturation (**E,F**), heart rate (**G,H**) and respiratory rate (**I,J**) is shown for each group. In all graphs, males are represented by filled symbols, and females by open symbols. For graphs displaying the nadir: Oxygen saturation: One-way ANOVA, $F_{(2,27)}$ = 0.1497, p=0.8617; heart rate: One-way ANOVA, $F_{(2,27)}$ = 2.115, p=0.1402; respiratory rate: One-way ANOVA, $F_{(2,27)}$ = 1.597, p=0.2211. For graphs displaying time to recover to 90% baseline: Oxygen saturation: One-way ANOVA, $F_{(2,27)}$ = 1.000, p=0.3811; heart rate: One-way ANOVA, F(2,27) = 0.7014, p=0.5047; respiratory rate: One-way ANOVA, F(2,27) = 1.216, p=0.3121.

| Table 1. Baseline cardiorespiratory values in rats prior to receiving 20 µg/kg fentanyl | Figure 1 Dose Response | | Figure 3 Pretreatment | | Figure 4 Reversal | |
|---|---|---|---|---|---|---|
| | Males | Females | Males | Females | Males | Females |
| Oxygen Saturation Baseline (%) | 94 ± 0.5 | 95 ± 0.4 | 96 ± 0.4 | 96 ± 0.4 | 95 ± 0.6 | 95 ± 0.3 |
| Heart Rate Baseline (beats per minute) | 445 ± 16 | 426 ± 13 | 462 ± 12 | 439 ± 17 | 436 ± 10 | 416 ± 13 |
| Respiratory Rate Baseline (breaths per minute) | 109 ± 6 | 95 ± 3 | 108 ± 5 | 114 ± 3 | 103 ± 4 | 102 ± 4 |

**Supplemental Table 1: Baseline cardiorespiratory values in rats prior to receiving 20 µg/kg fentanyl**. Values are mean ± SE. Data are from rats used in Figures 1, 3 and 4 and are separated by sex. There was no significant difference in any of the measured parameters between male and female rats. Oxygen saturation: one-way ANOVA, $F_{(5, 27)} = 0.6787$, p =0.6424; heart rate: one-way ANOVA, $F_{(5, 27)} = 1.078$, p=0.3944; respiratory rate: one-way ANOVA, $F_{(5, 27)} = 1.527$, p=0.2147.

| Table 2. Baseline cardiorespiratory values in rats prior to receiving 50 µg/kg fentanyl | Figure 1 Dose Response | | Figure S1 Pretreatment | | Figure S2 Reversal | |
|---|---|---|---|---|---|---|
| | Males | Females | Males | Females | Males | Females |
| Oxygen Saturation Baseline (%) | 94 ± 1 | 95 ± 1 | 94 ± 1 | 96 ± 1 | 94 ± 1 | 95.2 ± 1.1 |
| Heart Rate Baseline (beats per minute) | 408 ± 10 | 485 ± 13* | 450 ± 21 | 434 ± 14 | 421 ± 17 | 451 ± 33 |
| Respiratory Rate Baseline (breaths per minute) | 95 ± 2 | 110 ± 4 | 101 ± 5 | 98 ± 4 | 100 ± 1 | 105 ± 8 |

**Supplemental Table 2: Baseline cardiorespiratory values in male and female rats prior to receiving 50 µg/kg fentanyl**. Values are mean ± SE. Data are from rats used in Figures 1, S1 and S2 and are separated by sex. Female rats in Figure 1 had a significantly higher heart rate at baseline compared male rats in Figure 1. Oxygen saturation: one-way ANOVA, $F_{(5, 25)} = 1.161$, p=0.3558; heart rate: one-way ANOVA, $F_{(5, 25)} = 3.774$, p=0.0110, Tukey's post-hoc test † $p<0.05$ Figure 1 males vs Figure 1 females; respiratory rate: one-way ANOVA, $F_{(5, 25)} = 1.728$, p=0.1649.

| **Table 3. Nadir and recovery values in male and female rats that received 20 µg/kg fentanyl** | Figure 1 Dose Response | | Figure 3 Pretreatment | | Figure 4 Reversal | |
|---|---|---|---|---|---|---|
| | Males | Females | Males | Females | Males | Females |
| Oxygen Saturation Nadir (%) | 50 ± 4 | 50 ± 4 | 42 ± 6 | 38 ± 1 | 53 ± 6 | 45 ± 2 |
| Heart Rate Nadir (beats per minute) | 182 ± 26 | 144 ± 10 | 253 ± 32 | 177 ± 55 | 226 ± 45 | 146 ± 20 |
| Respiratory Rate Nadir (breaths per minute) | 59 ± 3 | 49 ± 4 | 67 ± 4 | 45 ± 12 | 66 ± 5 | 60 ± 4 |
| Oxygen Saturation Recovery (minutes) | 11 ± 1 | 10 ± 1 | 12 ± 2 | 14 ± 2 | 14 ± 3 | 11 ± 1 |
| Heart Rate Recovery (minutes) | 15 ± 13 | 16 ± 1 | 15 ± 3 | 13 ± 5 | 15 ± 3 | 18 ± 1 |
| Respiratory Rate Recovery (minutes) | 12 ± 2 | 14 ± 1 | 10 ± 1 | 17 ± 4 | 11 ± 1 | 15 ± 3 |

**Supplemental Table 3: Nadir and recovery values in male and female rats that received 20 µg/kg fentanyl**. Values are mean ± SE. Data are from rats used in Figures 1, 3 and 4 and are separated by sex. For nadir data: oxygen saturation: one-way ANOVA, $F_{(5, 27)} = 1.229$, p =0.3230; heart rate: one-way ANOVA, $F_{(5, 27)} = 1.844$, p=0.1379; respiratory rate: one-way ANOVA, $F_{(5, 27)} = 2.669$, p=0.0438. Despite the significant interaction for respiratory rate, no post hoc differences were detected. Sex differences in recovery times were also evaluated. oxygen saturation: one-way ANOVA, $F_{(5, 27)} = 0.7021$, p =0.6267; heart rate: one-way ANOVA, $F_{(5, 27)} = 0.4536$, p=0.8069; respiratory rate: one-way ANOVA, $F_{(5, 27)} = 1.100$, p=0.3832.

| **Table 4. Nadir and recovery values in male and female rats that received 50 µg/kg fentanyl** | Figure 1 Dose Response | | Figure S1 Pretreatment | | Figure S2 Reversal | |
|---|---|---|---|---|---|---|
| | Males | Females | Males | Females | Males | Females |
| Oxygen Saturation Nadir (%) | 30 ± 3 | 37 ± 2 | 41 ± 4 | 35 ± 6 | 40 ± 5 | 38 ± 4 |
| Heart Rate Nadir (beats per minute) | 181 ± 16 | 145 ± 20 | 159 ± 17 | 179 ± 37 | 135 ± 17 | 188 ± 33 |
| Respiratory Rate Nadir (breaths per minute) | 48 ± 6 | 41 ± 5 | 56 ± 10 | 55 ± 9 | 50 ± 8 | 61 ± 6 |
| Oxygen Saturation Recovery (minutes) | 31± 6 | 28 ± 7 | 30 ± 5 | 22 ± 6 | 29 ± 5 | 28 ± 6 |
| Heart Rate Recovery (minutes) | 23 ± 5 | 48 ± 1 | 43 ± 9 | 33 ± 5 | 35 ± 7 | 41 ± 1 |
| Respiratory Rate Recovery (minutes) | 16 ± 3 | 43 ± 1 †# | 26 ± 7 | 18 ± 4 | 13 ± 4 | 38 ± 3 |

**Supplemental Table 4: Nadir and recovery values in male and female rats that received 50 µg/kg fentanyl**. Values are mean ± SE. Data are from rats used in Figures 1, S1 and S2 and are separated by sex. For nadir data: oxygen saturation: one-way ANOVA, $F_{(5, 25)} = 1.384$, p =0.2639; heart rate: one-way ANOVA, $F_{(5, 25)} = 0.8879$, p=0.5039; respiratory rate: one-way ANOVA, $F_{(5, 25)} = 1.154$, p=0.3591. Sex differences in recovery times were also evaluated. oxygen saturation: one-way ANOVA, $F_{(5, 25)} = 0.2277$, p =0.9469; heart rate: one-way ANOVA, $F_{(5, 25)} = 1.591$, p=0.1992; respiratory rate: one-way ANOVA, $F_{(5, 25)} = 4.258$, p=0.0061. Tukey's post-hoc test † $p<0.05$ Figure 1 males vs Figure 1 females, # $p<0.05$ Figure 1 females vs Figure S2 males.
